# Dual regulation of the receptor-like kinase *BIR1* involves site-directed transcript cleavage and 5’-leader-mediated translational control

**DOI:** 10.64898/2026.01.16.699855

**Authors:** Irene Guzmán-Benito, Carmen Robinson, Livia Donaire, Lourdes Fernández-Calvino, José Manuel Franco-Zorrilla, Ruixia Niu, Guoyong Xu, Catharina Merchante, César Llave

## Abstract

In Arabidopsis, receptor-like kinase BRASSINOSTEROID INSENSITIVE1-ASSOCIATED RECEPTOR KINASE 1 (BAK1)*-*INTERACTING RECEPTOR-LIKE KINASE 1 (BIR1) is a negative regulator of plant immunity and cell death. BIR1 was earlier described as a target of epigenetic and post-transcriptional silencing. Degradome analysis mapped predominant mRNA cleavage sites at the 5’-untranslated leader region (site A) and the protein-coding sequence (sites B and C). Here, we identified another cleavage site (D) within the *BIR1* coding region and investigated the contribution of site-directed mRNA cleavage to *BIR1* regulation. Mutations at B, C, and D sites enhanced mRNA stability by impairing transcript cleavage, resulting in increased *BIR1* mRNA and protein accumulation. This regulation is disrupted in RNA silencing mutants, supporting a model of *cis*-directed small interfering RNA (siRNA)-mediated degradation. Cleavage events are highly localized, occurring at a limited number of sites that involve only a few siRNAs. Furthermore, our data reveal a repressive role for the 5’-leader in regulating *BIR1* translation, potentially mediated by upstream open reading frames (uORFs) and a long non-coding RNA (lncRNA) derived from the natural antisense *At4g39838* locus. This lncRNA is complementary to a TCA-rich repeat region encompassing cleavage site A, which may serve as a putative binding site. Together, these findings reveal a multilayered regulatory mechanism that integrates sRNA-mediated cleavage with translational control, with broader implications for the fine-tuning of stress-responsive gene expression during infection.

## Introduction

Plants have evolved intricate mechanisms to fine-tune the strength and duration of the immune responses, highlighting the importance of the immune attenuation to prevent self-damage from prolonged or excessive activation (Hou and Xu, 2025; Tang *et al*., 2025). Plant innate immunity comprises pathogen-associated molecular pattern (PAMP)-triggered immunity (PTI) and effector-triggered immunity (ETI) (Jones *et al*., 2024). PTI is mediated by cell surface pattern recognition receptors (PRRs), while ETI involves intracellular nucleotide-binding domain leucine-rich repeat receptors (NLRs) (Zhou and Zhang, 2020). NLRs are encoded by resistance (*R*) genes and sense pathogen-effector proteins or effector-induced manipulations of host proteins (Duxbury *et al*., 2021; Yu *et al*., 2021). PTI and ETI function synergistically and interdependently in plants and share common downstream responses (Jones and Dangl, 2006; Pruitt *et al*., 2021; Tian *et al*., 2021). Besides, ETI often triggers a form of programmed cell death called hypersensitive response (HR) (Coll *et al*., 2011).

PTI modulation involves multiple layers, including the formation of PRR-co-receptor complexes, regulation of signal initiation, cytoplasmic signal transduction, and transcriptional control (Couto and Zipfel, 2016; Mithoe and Menke, 2018; Withers and Dong, 2017). Post-translational mechanisms such as phosphorylation, ubiquitination, and lipid modifications play critical roles in modulating plant immune signaling. Phosphatases like PP2A and PP2C38 attenuate immune responses by regulating the activity of receptor-like kinases (RLKs) BRASSINOSTEROID INSENSITIVE1-ASSOCIATED RECEPTOR KINASE 1 (BAK1) and BOTRYTIS INDUCED KINASE1 (BIK1), respectively (Couto and Zipfel, 2016; Segonzac *et al*., 2014; Song *et al*., 2026). The ubiquitin-proteasome system controls PRR abundance, as seen in BAK1-mediated activation of E3 ubiquitin ligases (PUB) that target FLAGELLIN-SENSING 2 (FLS2) and BIK1 for degradation (Lu *et al*., 2011; Mithoe and Menke, 2018; Wang *et al*., 2018). Additional modifications, including glycosylation and acylation, further influence RLK stability and localization (Withers and Dong, 2017). Finally, MAPK cascades and WRKY transcription factors help balance early PTI signaling through feedback regulation of defense gene expression (Bi *et al*., 2018; Couto and Zipfel, 2016).

Similarly, NLR-mediated immune pathways are tightly regulated at multiple levels, including transcriptional, post-transcriptional, translational, and post-translational control. Among these, RNA silencing serves as a key checkpoint modulating both PTI and ETI responses in plants (Huang *et al*., 2016; Lopez-Marquez *et al*., 2023). RNA silencing is driven by two major classes of small RNAs (sRNAs): ∼21-nt microRNAs (miRNAs) and 21–24 nt small interfering RNAs (siRNAs) (Lopez-Gomollon and Baulcombe, 2022). Mounting evidence highlights the role of pathogen-responsive miRNAs and siRNAs in plant innate immunity (Boccara *et al*., 2014; Campo *et al*., 2013; He *et al*., 2008; Katiyar-Agarwal *et al*., 2007; Katiyar-Agarwal *et al*., 2006; Li *et al*., 2014; Li *et al*., 2010; Navarro *et al*., 2006; Navarro *et al*., 2008; Ouyang *et al*., 2014; Zhang *et al*., 2011). These sRNA networks exert broad post-transcriptional control over NLRs encoded by resistance genes through target RNA cleavage or translational repression, thereby preventing autoimmunity from excessive NLR accumulation (Boccara *et al*., 2014; Li *et al*., 2012; Shivaprasad *et al*., 2012; Yi and Richards, 2007; Zhai *et al*., 2011). RNA-directed DNA methylation (RdDM) contributes to immune regulation by silencing transposable elements and nearby defense genes under non-stress conditions (Dowen *et al*., 2012; Le *et al*., 2014; Lopez Sanchez *et al*., 2016). Upon pathogen attack, DNA demethylation can lift this repression, activating immune responses and restricting infection (Navarro *et al*., 2004; Yu *et al*., 2013).

The RLK BAK1-INTERACTING RECEPTOR-LIKE KINASE 1 (BIR1) exemplifies another immune regulator whose expression is controlled by both epigenetic and post-transcriptional silencing (Guzman-Benito *et al*., 2019). BIR1 is one of the four members of the BIR family, of which BIR1, BIR2 and BIR3 function to prevent inappropriate activation of BAK1-mediated PRR signaling. Under non-infectious conditions, BIR proteins keep BAK1 inactive through constitutive interaction, but upon pathogen detection, they dissociate, enabling BAK1 to associate with activated PRRs and trigger immune signaling (Gao *et al*., 2009; Halter *et al*., 2014; Imkampe *et al*., 2017). Notably, both *bir1-1* mutants and *BIR1* overexpression lines in Arabidopsis show autoimmune-like phenotypes (Gao *et al*., 2009; Guzman-Benito *et al*., 2019), emphasizing the importance of precise *BIR1* regulation. RdDM ensures that *BIR1* expression remains balanced under non-stress conditions (Guzman-Benito *et al*., 2019). However, upon infection, *BIR1* is transcriptionally induced, likely through a regulatory mechanism involving antagonistic salicylic acid (SA) and jasmonic acid (JA) interactions (Robinson *et al*., 2025). Under these conditions, post-transcriptional gene silencing (PTGS) reinforces epigenetic repression to prevent excessive accumulation. This idea is supported by elevated *BIR1* transcript levels in infected RNA silencing mutants and the accumulation of *BIR1*-derived siRNAs specifically in infected plants (Guzman-Benito *et al*., 2019). Degradome analysis further revealed increased cleavage of *BIR1* transcripts in infected tissues, indicating active transcript degradation (Guzman-Benito *et al*., 2019).

In this study, we identified *BIR1* as part of a broad set of plant genes whose mRNAs undergo selective degradation during viral infections but not under control conditions. Mutational analysis of degradome-derived 5’ cleavage sites showed that *BIR1* is post-transcriptionally regulated by sRNA-dependent, sequence-specific endonucleolytic cleavage in its coding region. Cleavage displays high specificity, being restricted to a few defined sites targeted by a limited repertoire of sRNAs. In addition, we identified several upstream open reading frames (uORFs) as well as a long non-coding RNA (lncRNA) with complementarity to the TCA-rich region of the *BIR1* 5’-leader, both of which contribute to the regulation of *BIR1* translation. Together, these findings provide new insights into the mechanisms governing *BIR1* expression.

## Materials and methods

### Plant material and virus inoculation

*Arabidopsis thaliana* and *Nicotiana benthamiana* plants were grown in controlled chambers under long-day conditions (16h day/8h night) at 22 and 24°C, respectively. Arabidopsi*s* mutants used in this study included *ago1-27, ago2-1*, *ago4-2*, *ago6-3* and *ago10-3* and the triple *dcl2-1 dcl3-1 dcl4-2* (provided James C. Carrington, The Donald Danforth Plant Center, MO, USA). T-DNA insertion Arabidopsis lines *SALK_097507C* and *SALK_093172C* were obtained from the Nottingham Arabidopsis Stock Centre (NASC). The *BIR1-mCherry* lines L6 and L9 under a dexamethasone (DEX)-inducible system were as described previously (Guzman-Benito *et al*., 2019). The same BIR1-mCherry system was used to generate BIR1-mCherry mutants containing nucleotide substitutions. Sequence mutations were introduced into the *BIR1* coding sequence by overlapping PCRs. These constructs were used for agroinfiltration assays and to transform Arabidopsis via floral dipping. Independent homozygous lines were selected in T3 for further experiments. Morphological phenotypes and gene expression after DEX treatments were examined in at least two independent transgenic lines. The *N. benthamiana* transgenic RNAi line NbDCL2/4i contains a chimeric hairpin construct targeting simultaneously *DCL2* and *DCL4* genes as described (Dadami *et al*., 2013) (provided by Kriton Kalantidis, Institute of Molecular Biology and Biotechnology, Crete, Greece).

The Arabidopsis *BIR1* genomic sequence (*At5g48380*) annotated in TAIR11 was PCR-amplified and cloned into the pGWB14 binary vector or the modified pYL156 vector, both carrying a C-terminal triple human influenza hemagglutinin (3xHA) tag. The 5’-leader sequence of *BIR1* was inserted upstream of the green fluorescence protein (GFP) or red fluorescence protein (RFP) coding regions into the pGWB405 and pGWB554 binary vectors, respectively. *N. benthamiana* plants were agro-inoculated with a tobacco rattle virus (TRV) infectious clone expressing green fluorescent protein (GFP) or *BIR1* derivatives under the replicase promoter of the pea early browning virus as described (Donaire *et al*., 2008). Three weeks after germination, Arabidopsis plants were mechanically inoculated with sap from upper, non-inoculated leaves of *N. benthamiana* plants. Sap was prepared by grinding infected tissue in 0.5M phosphate buffer (pH 7.0) at a 1:1 (W/V) ratio and applied onto the leaf surface using carborundum as an abrasive. Relative TRV levels in *N. benthamiana* plants were measured on upper non-inoculated leaves 5 days post-inoculation (dpi) using RT-qPCR.

### Transient expression in *N. benthamiana*

Transient expression was conducted in *N. benthamiana* plants grown under long day conditions (16h day/8h night at 25°C). Agroinfiltration of *N. benthamiana* leaves was performed as described previously (Johansen and Carrington, 2001). In co-infiltration assays, cultures containing the viral constructs were infiltrated at OD_600_ of 0.2, cultures harboring lncRNA or miRNA precursor constructs at OD_600_ of 0.4 and cultures carrying *BIR1* derivatives were infiltrated at OD_600_ of 0.4. The final concentration of bacteria was normalized within each experiment by varying the concentration of cells containing empty vector.

### RNA analysis and Quantitative RT-PCR assay

Total RNA was extracted using TRIzol reagent (Invitrogen). Denaturing gel blot hybridization of normalized total RNA was performed as previously described (Martinez-Priego *et al*., 2008). Radiolabeled DNA probes were synthesized by random priming with [α-³²P]-dCTP. One-step qRT-PCR was conducted using Brilliant III Ultra-Fast SYBR Green QRT-PCR Master Mix (Agilent Technologies) on a Rotor-Gene 6000 or Rotor-Gene Q real-time PCR system (Corbett/QIAGEN), following (Fernandez-Calvino *et al*., 2016). Relative gene expression was calculated using the ΔΔCt method with Rotor-Gene software. Expression was normalized to *CBP20* (*At5g44200*), which showed stable expression in both mock- and virus-infected samples. Primers used are listed in **Table S1**.

### Protein analysis

Leaf tissue was ground in liquid nitrogen and homogenized in extraction buffer (65 mM Tris-HCl, pH 8.0; 3% SDS; 1% β-mercaptoethanol; 10% glycerol). Samples were mixed with Laemmli buffer, heated at 95 °C for 5 min, and separated on 10% SDS-polyacrylamide gel electrophoresis (PAGE) gels. Proteins were transferred to P0.45 Polyvinylidene Fluoride (PVDF) blotting membranes (Amersham) and detected using HRP-conjugated secondary antibodies and chemiluminescent substrate (LiteAblot Plus).

### Small RNA sequencing, construction of degradome libraries and 5’ RACE

Young rosette leaves from virus-infected and corresponding mock-inoculated plants (10–12 per sample) were pooled at 8 dpi (TRV) or 14 dpi (TuMV) for degradome or small RNA sequencing. Infection was confirmed in each plant, and mock-inoculated controls were verified to be virus-free. Tissue was flash-frozen in liquid nitrogen and stored at –80 °C until use. Total RNA was extracted using TRIzol reagent (Invitrogen) or the RNeasy Plant Mini Kit (QIAGEN), and RNA quality was assessed with an Agilent 2100 Bioanalyzer. Small RNA libraries were prepared and sequenced as previously described (Guzman-Benito *et al*., 2019).

Arabidopsis degradome analysis was performed using two complementary high-throughput methods. First, differentially processed transcripts were identified via T7 RNA polymerase-mediated amplification of 5’-cleaved RNAs and hybridization to ATH1 microarrays (∼24,000 transcript variants from ∼22,500 genes), following the protocol by (Franco-Zorrilla *et al*., 2009). Three independent biological replicates and their respective negative controls (cDNA with RNA adapter present but not ligated) were hybridized independently. Genes with ≥2-fold enrichment and FDR ≤ 0.05 were considered significantly overrepresented. In parallel, degradome libraries were prepared from mock- and virus-infected plants for high-throughput sequencing using the PARE method, as described (Guzmán-Benito et al., 2019). Internal cleavage sites on predicted degradation targets were validated by 5’-RACE (Llave *et al*., 2002).

### Analyses of upstream Open Reading Frames (uORF)

Ribo-seq and RNA-seq datasets from Col-0 plants under control and ETI-induced conditions were obtained from Zhou *et al*. (2023) and analyzed following the procedures described therein. Reads were aligned to *BIR1*, and normalized read counts were quantified for the main open reading frame (mORF) and annotated upstream open reading frames (uORFs). Ratios of read counts were calculated between experimental conditions and between the mORF and uORFs.

### Data access

Arabidopsis genome annotations (TAIR11) were obtained from TAIR (www.arabidopsis.org). Degradome data from naïve Col-0 inflorescences (GSM280226) were previously published (German et al., 2008). Data from this study are available in the NCBI Gene Expression Omnibus (GEO) under the following accession numbers: GSE106322 (Microarray hybridization of degradome tags), GSM3019138-40 (Deep sequencing of degradome tags), GSM2808011-12, GSM3019141-42 (Deep sequencing of small RNAs).

## Results

### *BIR1* transcripts undergo endonucleolytic cleavage in virus-infected plants

In a previous study, degradome profiling of 5’ end signatures of uncapped and cleaved transcripts revealed specific cleavage hotspots within *BIR1* transcripts from TRV-infected plants, indicating that *BIR1* is a target of endonucleolytic processing during infections (Guzman-Benito *et al*., 2019). Notably, TRV infection was associated with prominent 5’ degradome signatures at nucleotide positions 156, 2,219, and 2,247 designated A, B, and C, respectively, which were clearly discernible from the background of low-abundance, non-specific degradation products (Guzman-Benito *et al*., 2019). To explore this further, we employed a genome-wide method previously described by Franco-Zorrilla et al. (2009) to identify candidate RNA silencing targets in mock-inoculated and TRV-infected Arabidopsis plants (Franco-Zorrilla *et al*., 2009). This approach uses parallel analysis of RNA ends (PARE) to generate libraries containing 3’ cleavage mRNA fragments, followed by Affymetrix ATH1 microarray hybridization of enriched cDNAs. Among the transcripts producing significant amounts of cleaved fragments relative to the hybridization controls, two genes (*At2g35750* and *At5g27860*) showing the highest fold enrichment under both conditions were selected for further analysis as RNA silencing targets (**Table S2**). RNA-seq-based degradome profiling, together with gene-specific 5’-RACE, confirmed specific and recurrent cleavage at defined sites in both transcripts (**Figure 1A**). In each case, the 5’-RACE products precisely matched the cleavage positions identified by the degradome data. Moreover, relative to wild-type plants (WT, Col-0), transcript levels of both genes were significantly increased in Arabidopsis mutants defective in slicer-competent AGO proteins (*ago1-27* and *ago2-1*), but not in mutants affecting RdDM-associated, non-slicing AGOs (*ago4-2*, *ago6-3*) (**Figure 1B**) (Vaucheret, 2008). Accumulation of *At2g35750*, but not *At5g27860*, transcripts increased in the *ago10-3* mutant allele. This suggests that *At2g35750* regulation also involves AGO10, which functions as a miRNA decoy in Arabidopsis (Yu *et al*., 2017). These results confirm that this approach reliably identifies *bona fide* targets of post-transcriptional RNA silencing. We then identified 47 and 98 genes with significantly enriched 3’ RNA cleavage products in mock-inoculated and TRV-infected plants, respectively (**Table S2**). As expected, we found that *BIR1* transcripts also accumulated significantly more 3’ degradation products in TRV-infected plants (*p* < 0.05) than in mock-treated controls, confirming that *BIR1* transcripts were selectively cleaved during viral infection (**Figure 1C**, **Table S2**). Using gene-specific 5’-RACE, we identified an additional cleavage site at position 1,817 (designated D; **Figure 1D**), distinct from those found by degradome sequencing (Guzman-Benito *et al*., 2019). This finding further supports the selective degradation of *BIR1* transcripts in response to TRV infection. None of the experimentally identified cleavage sites in *BIR1* matched annotated miRNAs in the miRBase database.

**Figure 1.**
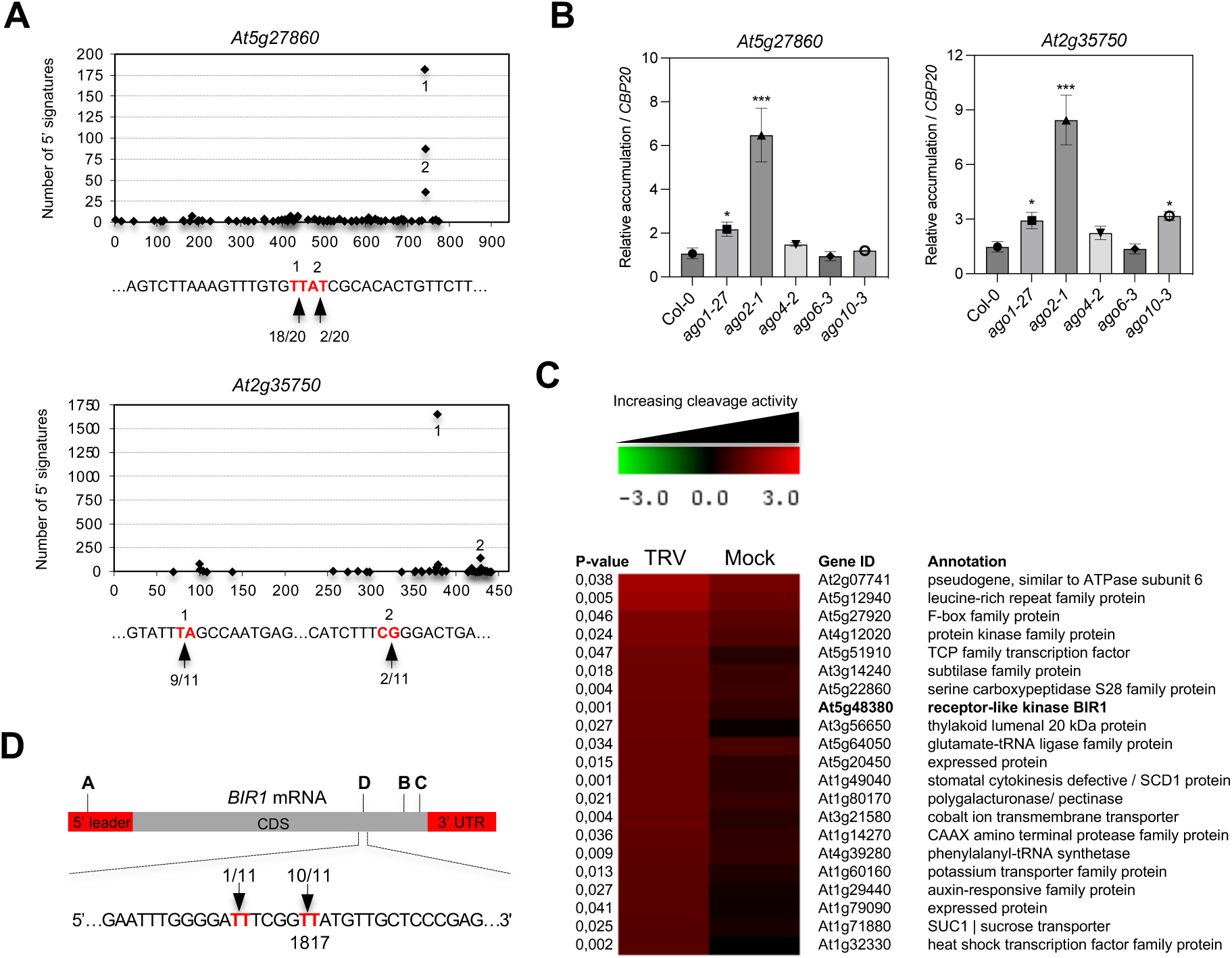
Genome-wide identification of degradation RNA targets using a Parallel-analysis of cDNA Ends (PARE)-based strategy. **(A)** Target plots derived from degradome sequencing showing 5’ cleavage signatures along the *At2g35750* and *At5g27860* mRNAs. Prominent peaks indicate preferential cleavage sites. Cleavage positions validated by 5’-RACE are shown below the plots and labeled with the corresponding numbers; clone counts indicate relative cleavage frequency. **(B)** RT-qPCR analysis of *At2g35750* and *At5g27860* transcript levels in wild-type plants (Col-0), slicer-competent ARGONAUTE (AGO) mutants (*ago1-27* and *ago2-1*), and RNA-Directed DNA Methylation (RdDM)-associated, non-slicing AGO mutants (*ago4-2*, *ago6-3*) ant the miRNA-related *ago10-3* mutant. Transcript levels were normalized to the *CBP20* internal control. Data represent mean ± SD (n=3) and were analyzed by one-way ANOVA followed by Tukey’s multiple-comparison test. **(C)** Heatmap showing Arabidopsis genes with the highest levels of differential RNA degradation between mock- and tobacco rattle virus (TRV)-infected plants, identified by PARE analysis followed by Affymetrix ATH1 microarray profiling. The color scale represents log₂(TRV/Mock) ratios; corresponding p-values (F-test, df = 1, 4) are indicated. *BIR1* (*At5g48380*) is highlighted in bold. **(D)** *BIR1* transcript cleavage sites identified by 5’-RACE in TRV-infected plants. Clone counts indicate cleavage frequency. Site D marks the most frequently detected cleavage position; additional cleavage sites (A-C) are also shown (Guzman-Benito *et al*., 2019). No 5’-RACE products were detected in mock-inoculated controls.

### Mutations within the predicted cleavage sites enhance *BIR1* expression

To determine whether cleavage of *BIR1* transcripts is site-specific, we first generated a *BIR1* variant carrying synonymous nucleotide substitutions at the three predicted cleavage sites within the *BIR1* open-reading frame. These mutations altered the RNA sequence at positions B, C, and D without affecting the encoded protein (**Figure 2A**, top). mCherry-tagged versions of the *BIR1* wild-type (WT) and the triple mutant (mBCD), including the 5’ untranslated leader region, were conditionally expressed in Arabidopsis using a transgenic DEX-inducible system described earlier (Guzman-Benito *et al*., 2019). Samples from two independent lines were collected after treatment with 30 µM DEX. RT-qPCR showed higher accumulation of mutated transcripts (BIR1 mBCD L3 and L7) compared to WT (BIR1 WT L6 and L9), indicating that these mutations confer cleavage resistance and thereby increase mRNA stability (**Figure 2B** and **Supplemental Figure 1A**). Consistently, BIR1 protein translated from mBCD transcripts in DEX-inducible L3 and L7 lines was more abundant than protein from WT transcripts in L6 and L9 transgenic lines (**Figure 2C**). Both BIR1-mCherry WT and mBCD proteins localized to the plasma membrane, ruling out altered subcellular localization as a cause of increased mBCD accumulation (**Figure 2D**).

**Figure 2.**
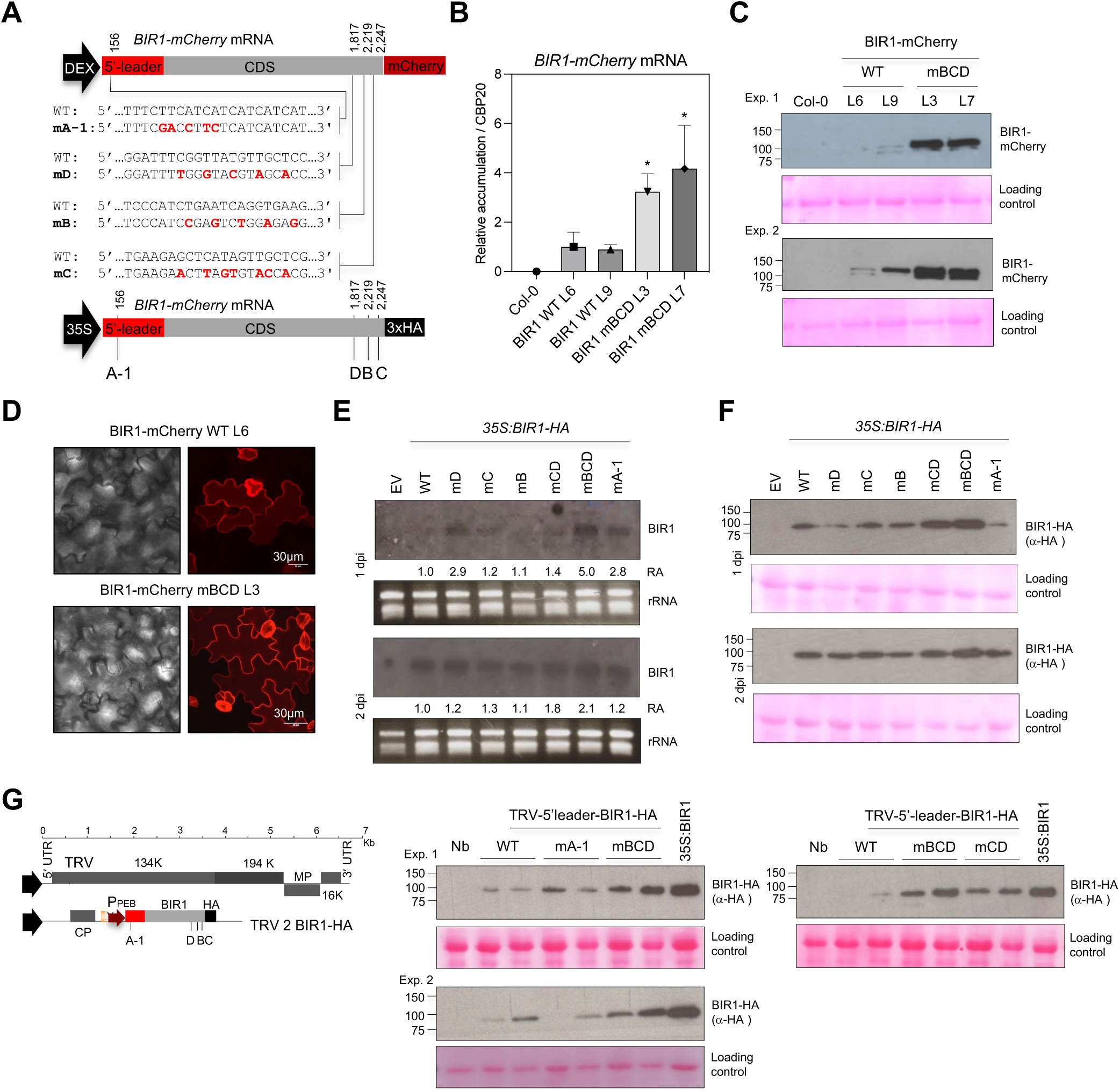
Effect of mutations at B, C and D cleavage sites within the protein-coding region on *BIR1* expression. (**A**) Top, schematic of DEX-inducible constructs expressing mCherry-tagged wild-type (WT) or mutant (m) *BIR1*. Mutations at cleavage sites A-D (nucleotide positions indicated) mutations are shown in red; sites B-D were altered by synonymous substitutions. Bottom, schematic of 35S-driven constructs expressing HA-tagged BIR1 variants. (**B**) RT-qPCR of *BIR1-mCherry* transcript levels in DEX-treated (16 h, 30 µM) Arabidopsis lines expressing WT (L6, L9) or mutated mBCD (L3, L7) *BIR1*. Transcript levels were normalized to the *CBP20* internal control. Data represent mean ± SD (n=3) and were analyzed by one-way ANOVA followed by Tukey’s multiple-comparison test. (**C**) Western blot of mCherry-tagged BIR1 proteins in DEX-treated (4 days) transgenic lines expressing WT (L6, L9) and mutated mBCD (L3, L7) BIR1-mCherry. Non-transgenic Col-0 served as a negative control. Results from two independent experiments (Exp. 1, Exp. 2) are shown. (**D**) Confocal images of mCherry fluorescence in DEX-treated (4 days) transgenic lines expressing WT (L6) and mBCD (L3) BIR1-mCherry. Scale bar = 30 µm. Imaging used a Leica TCS SP5 confocal microscope (561 nm excitation). (**E**) RNA blot of WT and mutant *BIR1* (mA-1, mB, mC, mD, mBC, mBCD) transcripts from 35S-driven constructs (top, A), sampled at 1- and 2-days post-infiltration (dpi). Blots were hybridized with *BIR1* cDNA probes and normalized to ethidium bromide staining. Relative accumulation (RA) shown with WT set to 1.0. Empty vector (EV) is the negative control. (**F**) Western blot of HA-tagged WT and mutated BIR1 proteins (same constructs as in B), collected at 1 and 2 dpi. (**G**) Left, schematic of tobacco rattle virus (TRV)-based constructs expressing HA-tagged BIR1 variants from the pea early browning virus (PEBV) replicase promoter in pTRV2. Transcription of pTRV1 and pTRV2 is 35S-driven. Cleavage sites A1-D are marked. Right, Western blot of HA-tagged WT and mutated BIR1 (mBCD, mA-1) from TRV-infected *Nicotiana benthamiana* leaves, sampled at 5 days post-inoculation (dpi). A 35S-driven BIR1-HA construct served as control. All immunoblots used anti-HA antibodies; Ponceau staining shows protein loading. Protein standards (KDa) are indicated. All *BIR1* constructs shown comprise the 5’ untranslated leader region followed by the *BIR1* coding sequence (CDS). Experiments were independently repeated with consistent results.

To determine the effects of single cleavage sites for *BIR1* mRNA regulation, we generated a panel of *BIR1* constructs carrying the silent mutations at each of the B, C, or D positions (**Figure 2A**, bottom). Both the WT and mutated constructs included the 5’-leader and a C-terminal 3x HA tag, and were transiently expressed in *N. benthamiana* under the 35S promoter. *BIR1* mRNA and protein levels were assessed at 1- and 2-days post-infiltration (dpi) by Northern and Western blot, respectively. RNA blots revealed that single mutations (mB, mC, or mD) had little or no effect on *BIR1* transcript accumulation (**Figure 2E**). In contrast, mutation of all three *BIR1* cleavage sites led to increased transcript accumulation relative to WT, consistent with reduced mRNA degradation and elevated expression (**Figure 2E**). Accordingly, BIR1 proteins translated from cleavage-resistant mBCD transcripts were also more abundant than those from WT transcripts or transcripts with single mutations (**Figure 2F**). These findings also confirmed that the observed effect was not tag-dependent.

To rule out agroinfiltration-related artifacts affecting *BIR1* expression, we further evaluated the impact of mBCD mutations using a viral-expression system. HA-tagged BIR1 WT and mBCD constructs, including their native 5’-leader, were cloned into the pTRV2 vector and co-delivered with pTRV1 into *N. benthamiana* via Agrobacterium infiltration (**Figure 2G**, left). *BIR1* expression was then analyzed in upper, non-infiltrated, systemically-infected leaves. As observed previously in the agroinfiltration assay, Western blot analysis showed higher levels of BIR1-HA protein derived from mBCD transcripts compared to WT in these leaves (**Figure 2G**, right). TRV RNA levels were comparable across WT, mBCD, and green fluorescence protein (GFP) controls in systemic leaves, indicating that the differences in *BIR1* expression were not due to variations in virus accumulation (**Supplemental Figure 1B**). As a control, a TRV-derived BIR1 construct carrying mutations at cleavage site A within the 5’-leader (mA-1) accumulated BIR1 protein at levels similar to the WT (**Figure 2G**). Collectively, our results across several expression systems and plant species confirm that *BIR1* transcripts undergo site-specific cleavage within the protein-coding region, which is critical for *BIR1* mRNA regulation.

### Site-directed cleavage at positions B, C, and D involves sRNA-mediated silencing pathways

Previous studies suggest that post-transcriptional gene silencing regulates *BIR1* expression, particularly during infection, when *BIR1*-derived siRNAs accumulate (Guzman-Benito *et al*., 2019). To confirm this, we generated Arabidopsis transgenic lines expressing *BIR1-mCherry* under a DEX-inducible promoter in the *dcl2 dcl3 dcl4* (herein *dcl234*) mutant background defective in sRNA biosynthesis and RNA silencing amplification. RT-qPCR analysis showed higher levels of *BIR1-mCherry* WT transcripts in two independent *dcl234* lines compared with the BIR1 L9 control (**Figure 3A**), supporting the idea that site-directed cleavage of *BIR1* transcripts involves the activity of RNA silencing-associated sRNAs. To further investigate this possibility, we agro-injected DEX-inducible, mCherry-tagged *BIR1* constructs in *N. benthamiana* NbDCL2/4i plants (**Figure 2A**, top). Western blot analysis showed higher accumulation of the mBCD protein than WT in WT *N. benthamiana*, whereas this difference was abolished in NbDCL2/4i plants, in which sRNA-mediated cleavage is impaired (**Figure 3B**). These findings suggest that cleavage at sites B, C, and D depends on DCL2/4-mediated sRNA pathways. As an additional control, expression of *BIR1* transcripts carrying the mA-1 mutation in the 5′ leader resulted in comparable protein levels across both genetic backgrounds (**Figure 3C**), indicating that site A is not regulated in the same way. Together, these results support a model in which *BIR1* is post-transcriptionally regulated via sRNA-guided cleavage at sites B, C, and D.

**Figure 3.**
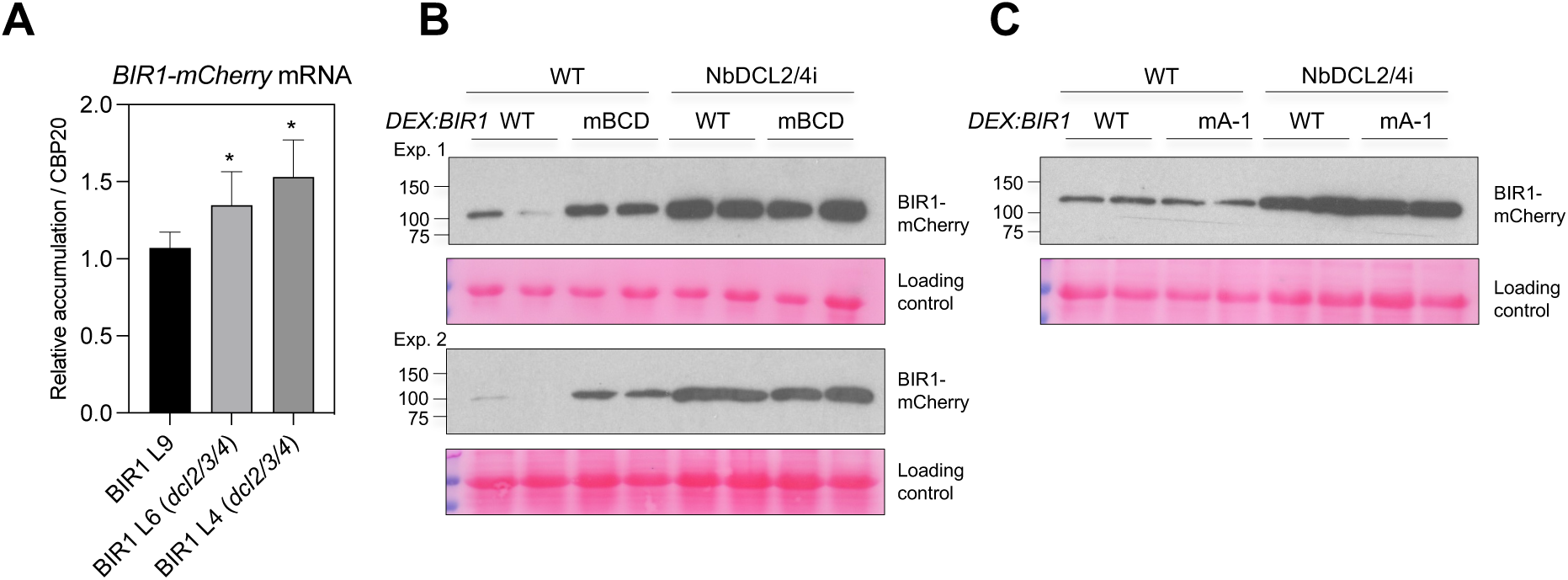
Cleavage at sites B, C, and D occurs through mechanisms involving sRNA-mediated silencing. (**A**) RT-qPCR of *BIR1-mCherry* transcript levels in transgenic plants of Col-0 (L9) and *dcl2 dcl3 dcl4* (*dcl234*; L4, L6) mutant genotypes expressing WT *BIR1-mCherry*, 24 h after DEX treatment. Transcript levels were normalized to the *CBP20* internal control. Data represent mean ± SD (n=5) and were analyzed by one-way ANOVA followed by Tukey’s multiple-comparison test. (**B**) Western blot of mCherry-tagged proteins WT and mBCD BIR1 in *Nicotiana benthamiana* WT and NbDCL2/4i leaves, 1-day post-infiltration (dpi) after DEX induction (30 µM). Results from two independent experiments (Exp. 1, Exp. 2) are shown. (**C**) Western blot of mCherry-tagged WT and mA-1 BIR1 in *N. benthamiana* WT and NbDCL2/4i, as in (B). Immunoblots used anti-mCherry antibodies; Ponceau staining shows loading controls. Protein standards (KDa) are indicated. Two samples per construct/genotype are shown. 35S-driven *BIR1* constructs shown comprise the 5’ untranslated leader region followed by the *BIR1* coding sequence (CDS). All experiments were repeated with similar results.

### *BIR1* overexpression compromises its post-transcriptional regulation

We further explored whether site-directed transcript cleavage alone was sufficient to contain the expression of *BIR1* in the absence of transcriptional regulation. To test the hypothesis, we used the DEX-inducible system to overexpress the mBCD mutated version of *BIR1-mCherry* in independent transgenic lines (BIR1 mBCD L3 and L7). Overexpression of *BIR1* induces cell death phenotypes and triggers transcriptional responses that greatly overlap with those of canonical PTI and ETI (Guzman-Benito *et al*., 2019). RT-qPCR showed that daily DEX applications for 10 days resulted in elevated transcript levels of both WT and mBCD compared to mock-treated controls (**Figure 4A**). Western blot analysis revealed slightly higher levels of BIR1 protein derived from mBCD transcripts compared with the WT at this time point (**Figure 4B**). Overexpression of *BIR1*-mBCD led to developmental and morphological defects characteristic of *BIR1* overexpression, appearing between 10 and 12 days after induction, similar to what was observed with the WT version. Interestingly, a slightly higher proportion of BIR1-mBCD transgenic plants exhibited these morphological defects following DEX treatments, suggesting that increased protein levels resulting from the loss of site-directed cleavage in the mBCD mutant may enhance the manifestation of cell death-associated morphologies (**Figure 4C**). Cell death and induction of the defense marker genes *PATHOGENESIS-RELATED 1* (*PR1)* and *PR4*, reported upon *BIR1* WT overexpression (Guzman-Benito *et al*., 2019), were also observed when mBCD *BIR1* was expressed via the TRV viral vector in Arabidopsis (**Figure 4D,E**) (Guzman-Benito *et al*., 2019). These findings indicate that high-level transcriptional activation of *BIR1* overrides the effects of site-directed transcript regulation. Thus, while sequence-specific cleavage normally constrains *BIR1* expression under physiological conditions, its regulatory impact is attenuated when *BIR1* is strongly overexpressed, possibly due to saturation or reduced efficiency of the RNA silencing machinery.

**Figure 4.**
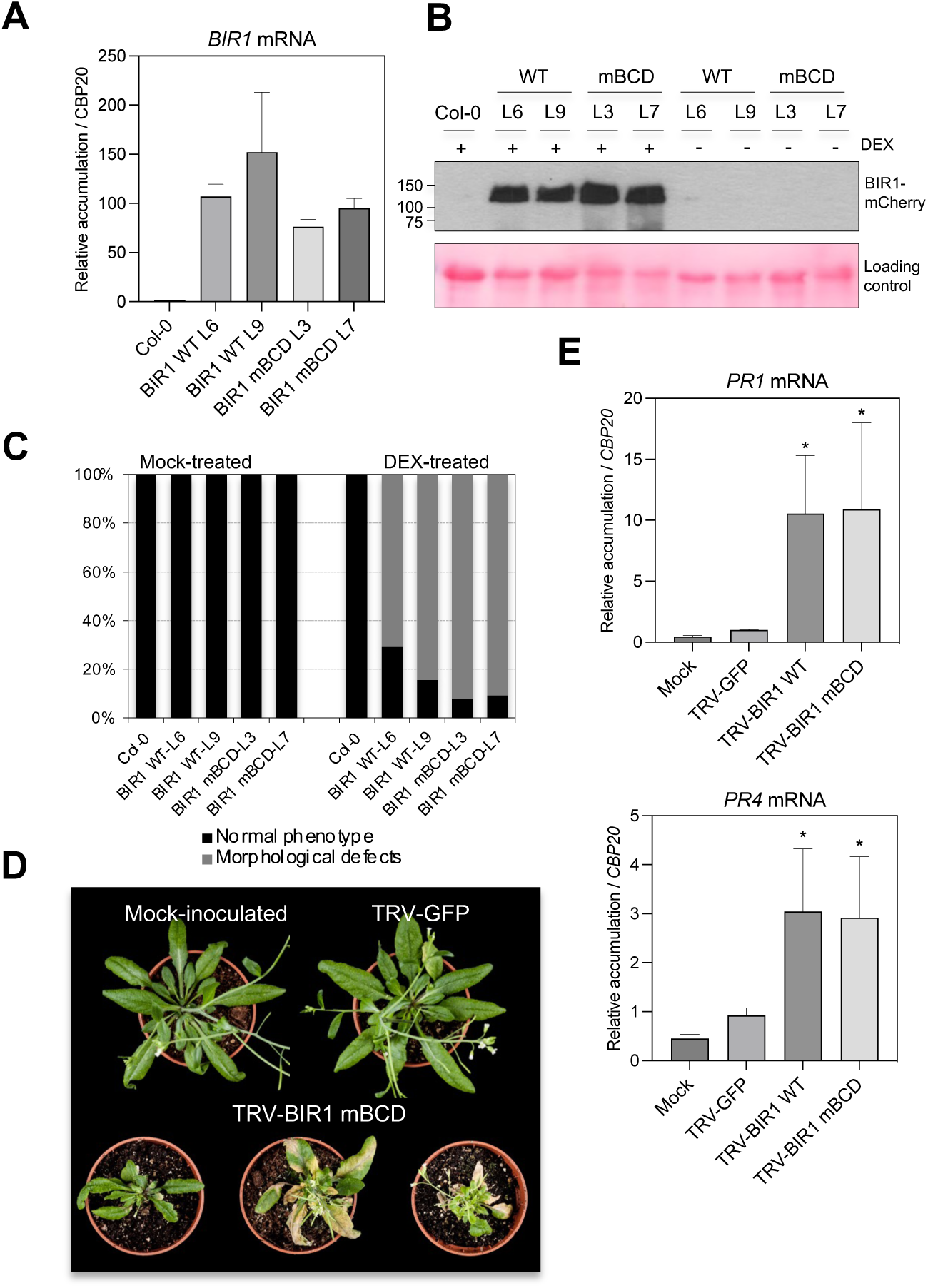
*BIR1* overexpression interferes with site-directed transcript regulation. (**A**) RT-qPCR analysis of *BIR1-mCherry* transcripts in DEX-inducible Arabidopsis lines expressing wild-type (WT; L6, L9) or mutant (mBCD; L3, L7) BIR1. Plants were treated with 30 µM DEX (+) or H₂O (-) for 10 days. (**B**) Western blot of mCherry-tagged WT and mBCD BIR1 proteins in the same lines as in (A) after 10 days of DEX or H₂O treatment. Non-transgenic Col-0 served as a negative control. Immunoblots used anti-mCherry antibodies; Ponceau staining shows loading controls. (**C**) Percentage of plants with normal or altered morphology in WT (L6, L9) and mBCD (L3, L7) Arabidopsis lines after DEX treatment. Non-transgenic Col-0 and H₂O-treated plants were used as controls. (**D**) Morphology of soil-grown plants mock-inoculated, infected with tobacco rattle virus (TRV)-green fluorescence protein (GFP), or TRV expressing BIR1 mBCD. Images taken at 14 dpi. (**E**) RT-qPCR analysis of *PATHOGENESIS-RELATED 1* (*PR1*), and *PR4* transcripts in plants from (D). Mock-inoculated, and TRV-GFP-infected plants served as controls. Transcript levels were normalized to the *CBP20* internal control. Data represent mean ± SD (n=5) and were analyzed by one-way ANOVA followed by Tukey’s multiple-comparison test. *p < 0.05. *BIR1* constructs shown comprise the 5’ untranslated leader region followed by the *BIR1* coding sequence (CDS). All experiments were repeated with consistent results.

### *BIR1* translation is influenced by regulatory elements within its 5’-leader

The 5’-leader of plant mRNAs is instrumental in controlling mRNA stability, translation efficiency, and responsiveness to developmental or environmental cues (Hardy and Balcerowicz, 2024; Srivastava *et al*., 2018). Thus, we examined the role of the 5’-leader in *BIR1* translation by expressing several 35S-driven *BIR1*-coding constructs, with or without the 5’-leader sequence, using agroinfiltration in *N. benthamiana*. Constructs containing the 5’-leader produced lower levels of BIR1 protein compared to those without it, suggesting a repressive effect (**Figure 5B**). To test this further, we designed chimeric constructs expressing HA-tagged eGFP from transcripts with or without the *BIR1* 5’ leader. In both 35S-driven expression and TRV-mediated delivery in *N. benthamiana*, transcripts with the 5’-leader led to reduced GFP protein accumulation (**Figure 5C, D**). These results suggest that the 5’-leader region contains regulatory elements that negatively modulate translation. Noteworthy, when *BIR1* constructs lacking the 5’-leader were expressed via agroinfiltration (**Figure 5B**) or delivered via TRV (**Figure 5E**) in *N. benthamiana*, differences in WT and mBCD protein accumulation were attenuated, suggesting that site-specific cleavage, and the resulting differences in expression, are also influenced by elements located within the 5’-leader.

**Figure 5.**
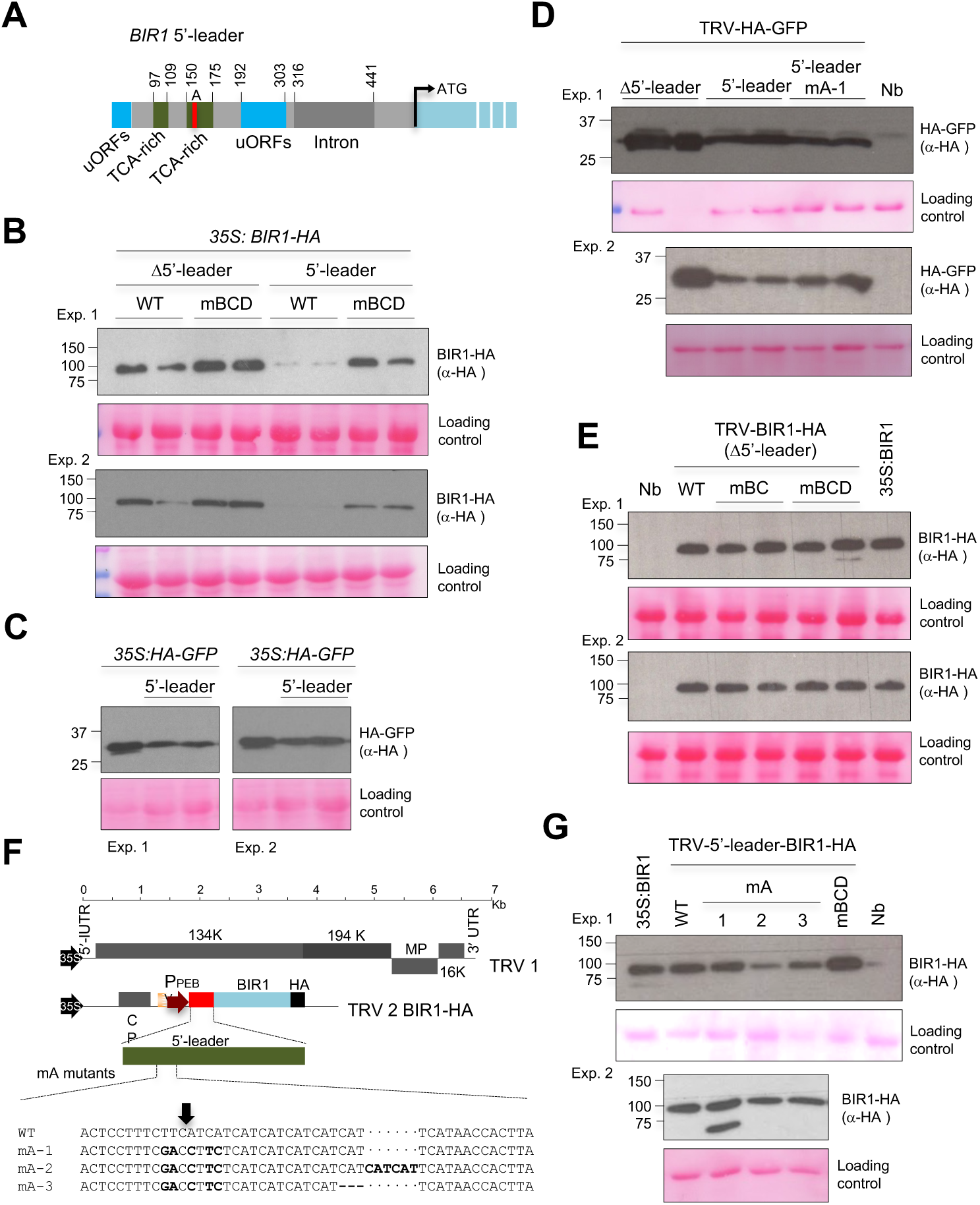
Effects of cleavage site mutations within the TCA-rich 5’ leader on *BIR1* expression. **(A)** Schematic of *cis*-regulatory elements within the 5’-leader of *BIR1*. The location and coordinates of upstream open reading frames (uORF), TCA-rich repeats and intron are shown. The canonical ATG start codon is shown. Cleavage site A is indicated in red. (**B**) Western blot comparing HA-tagged BIR1 expression with or without the 5’-leader, using 35S-driven WT and mBCD BIR1 constructs in the transient assay at 2 dpi. **(C)** Western blot comparing HA-tagged GFP expression with or without the 5’-leader using 35S-drive constructs in the transient assay at 2 dpi. **(D**) Western blot comparing HA-tagged GFP expression with or without the 5’-leader (WT and mutated mA-1) via TRV in *Nicotiana benthamiana*, sampled at 5 dpi. Results from two independent experiments (Exp. 1 and 2) are shown. The asterisk indicates a 4x diluted sample. (**E**) Western blot of HA-tagged WT and mutated BIR1 (mBC, mBCD) expressed from TRV constructs lacking the 5’-leader, sampled at 5 dpi. Results from two independent experiments (Exp. 1 and 2) are shown. (**F**) Schematic of tobacco rattle virus (TRV)-based constructs expressing HA-tagged wild-type (WT) or mutated *BIR1* (mA variants). Constructs include the *BIR1* 5’-leader and coding sequence (CDS), driven by the pea early browning virus (PEBV) replicase promoter in pTRV2. Transcription of pTRV1 and pTRV2 is 35S-driven. Cleavage sites A-D are marked; synonymous mutations are shown in **Figure 2A**. The major A site at position 156 (identified by degradome sequencing) is marked with a thick arrow. (**G**) Western blot of HA-tagged WT and mA mutant BIR1 proteins (mA-1, mA-2, mA-3) from TRV-infected *N. benthamiana* leaves (constructs from A), sampled at 5 dpi. A 35S-driven BIR1-HA construct served as control. Results from two independent experiments (Exp. 1 and 2) are shown. All immunoblots used anti-HA antibodies; Ponceau staining shows protein loading. Protein standards (KDa) are indicated. Experiments were independently repeated with consistent results.

### The *BIR1* leader contains multiple translated uORFs

We next examined the role of regulatory elements within the 5’-leader sequence of *BIR1* in modulating its expression. uORFs, which are small ORFs located within the 5’-leader sequences upstream of the main ORF (mORF), can act as molecular switches, activating or mostly repressing mORF translation in response to developmental and environmental cues (Nazarian-Firouzabadi *et al*., 2025; Yuan *et al*., 2024). According to the TAIR11 transcript annotation and uORFlight database (Niu *et al*., 2020), *BIR1* harbors six uORFs in its 5’-leader (**Figure 6**). In TAIR11, the transcription start site (TSS) is defined using the longest annotated 5′-end (Zhu *et al*., 2024). However, inspection of RNA-seq read coverage revealed that transcripts initiating from this longest 5′-end are rare. Consequently, the first three annotated uORFs are present only in low-abundance alternative TSS isoforms, whereas the majority of *BIR1* transcripts lack these three uORFs (**Figure 6**).

**Figure 6.**
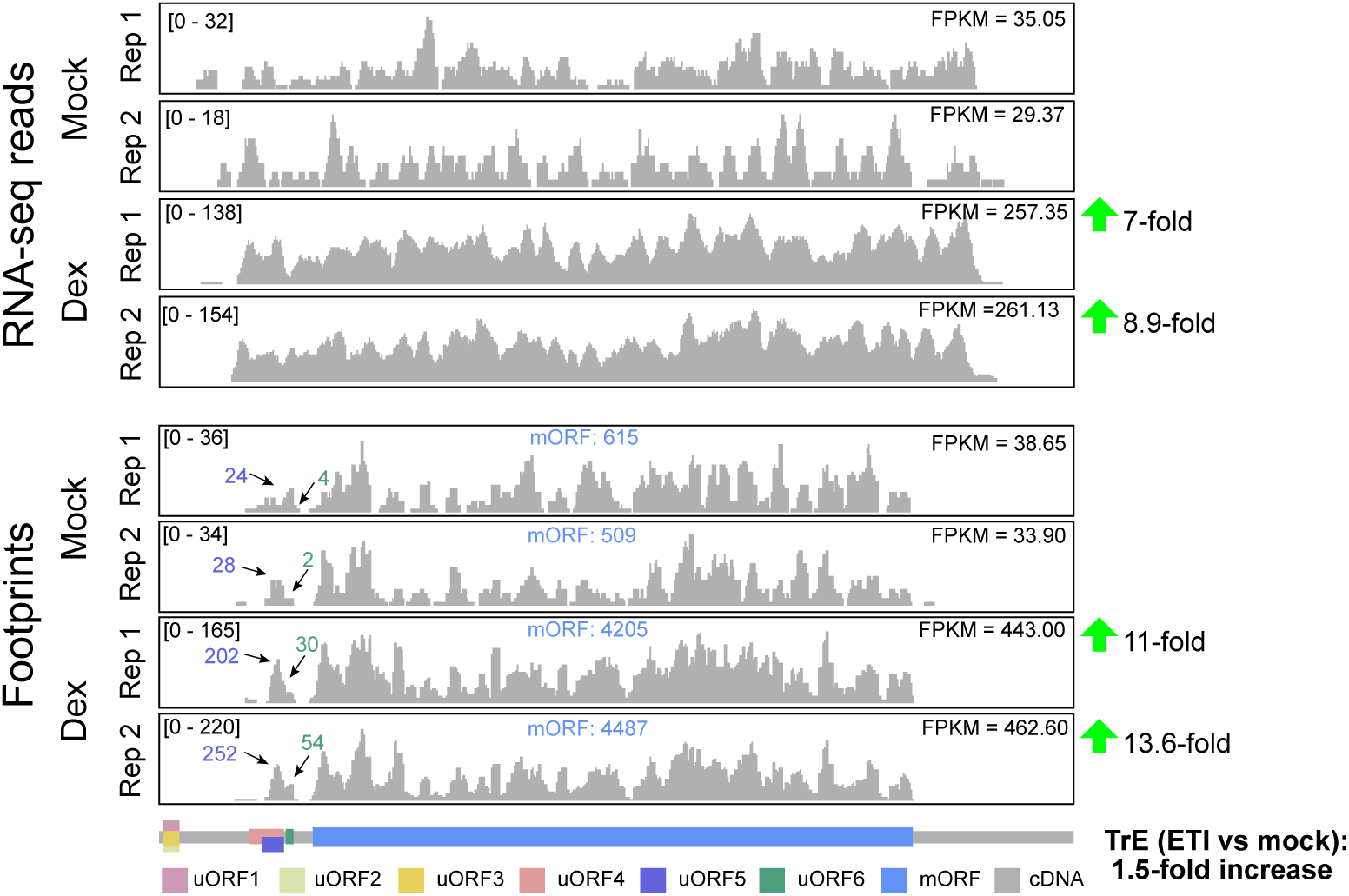
Translational control of *BIR1* expression involved multiple upstream open reading frames. Normalized distributions of RNA-seq and Ribo-seq reads along the *BIR1* transcript in Col-0 plants expressing a dexamethasone (Dex)-inducible AvrRpt2 effector from *Pseudomonas syringae* under mock and ETI-induced conditions. Transcript expression levels are given as Fragments Per Kilobase of transcript per Million mapped reads (FPKM). The grey bar represents the *BIR1* cDNA, with color-coded boxes indicating the six annotated upstream open reading frames (uORFs) and the main open reading frame (mORF). Numbers above the read profiles indicate read counts for uORF5, uORF6, and the mORF. Numbers following the green arrows denote fold changes in transcript and ribosome footprint abundance upon DEX treatment. The fold change in translational efficiency (TrE), representing the ratio of Ribo-seq to RNA-seq signals, is shown in bold. Data from (Zhou *et al*., 2023).

Therefore, our subsequent analyses focused on the three uORFs that are consistently present in the predominant transcript isoforms (uORFs 4, 5, and 6). To investigate whether these uORFs are translated and contribute to the regulation of *BIR1* expression, we analyzed publicly available Ribo-seq data from Arabidopsis Col-0 plants (Zhou *et al*., 2023). Normalized Ribo-seq and RNA-seq reads mapped along the *BIR1* transcript revealed ribosome footprints at all three uORFs, indicating that these *BIR1* uORFs are recognized by translating ribosomes (**Figure 6**). Noteworthy, uORFs translation also occurs under ETI-induced conditions, as observed in Arabidopsis plants expressing a DEX-inducible version of the AvrRpt2, an effector protein secreted by pathogenic *Pseudomonas syringae* (**Figure 6**). Consistent with previous reports (Guzman-Benito *et al*., 2019), ETI induction resulted in a marked increase in both *BIR1* mRNA abundance (∼8-fold) and translation (∼12-fold). This means that ETI enhanced the translation efficiency (TrE) of *BIR1* by approximately 1.5-fold (**Figure 6**). To assess whether this increase in translation could be attributed to altered translation dynamics of the uORFs, we compared the ratios of total Ribo-seq footprint reads in the mORF versus the uORFs, as well as between uORF5 and uORF6, under both mock and DEX-induced conditions. These analyses revealed no significant differences that could account for the observed increase in mORF TrE (**Figure 6**). However, given that these uORFs, particularly uORFs 5 and 6, are recognized by ribosomes and translated, they are likely to play a significant role in modulating *BIR1* translation.

### A long non-coding RNA (lncRNA) modulates *BIR1* translation

LncRNAs can regulate translation by base pairing with the 5’ unstranslated region (5’ UTR) of a target gene (Chekanova, 2021; Jampala *et al*., 2021). Accordingly, we searched the Arabidopsis genome for non-protein-coding loci and identified the *At4g39838* locus, which share partial sequence complementarity to the TCA-enriched region of the 5’-leader of *BIR1.* We hypothesized that this locus functions as a regulatory lncRNA, with the TCA repeats serving as its functional binding site. To test it, we first genotyped two Arabidopsis T-DNA insertion lines (SALK_093172C and SALK_097507C) targeting the overlapping *At4g39838*/*At4g39840* region. Both insertions were located upstream of the predicted lncRNA-binding sequence (**Supplemental Figure 2A**). RT-PCR showed that *At4g39838* transcripts were only slightly reduced in the homozygous mutants compared to the WT genotype, suggesting that *At4g39838*-related regulatory activity may still be retained in these mutants (**Supplemental Figure 2B**).

Alternatively, we generated two *At4g39838* constructs containing the predicted lncRNA sequence complementary to the *BIR1* 5’-leader, both driven by the constitutive 35S promoter (**Figure 7A**). These constructs were co-expressed with a 35S-driven HA-tagged *BIR1,* including the 5’-leader, using agroinfiltration in *N. benthamiana*. Strikingly, both *At4g39838*-derived lncRNA transcripts caused a strong decrease in BIR1 protein accumulation compared to the empty vector (**Figure 7B, D**). As expected, this effect was greatly reduced when a *BIR1* construct lacking the 5’-leader was used, indicating that this region was required for *At4g39838*-mediated repression of *BIR1* expression (**Figure 7B**). Moreover, co-expression of unrelated lncRNA harboring miRNA precursors (miR399, miR158, miR171) had no effect on BIR1 protein levels, regardless of the presence of the 5’-leader, confirming that the *At4g39838*-derived lncRNA operates specifically on regulatory elements within the 5’-leader of *BIR1* transcripts (**Figure 7C**). Together, these results demonstrate that the *At4g39838*-derived lncRNA exerts a sequence-specific, 5’-leader-dependent regulatory effect on *BIR1* expression, and identify the TCA-rich region as a potential regulatory element within the 5’-leader.

**Figure 7.**
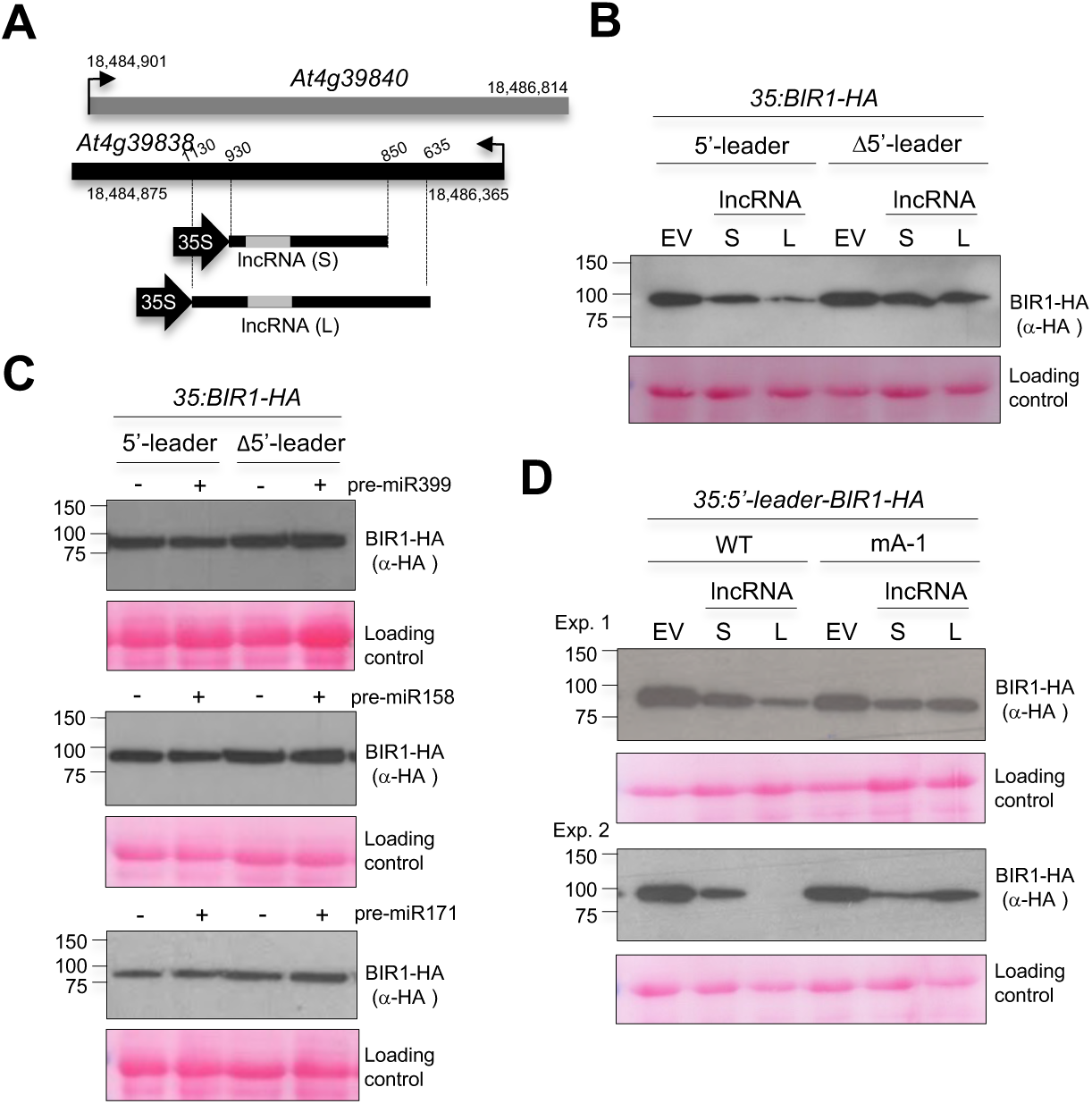
Expression of a *At4g39838*-derived lncRNA reduces BIR1 protein accumulation in Agrobacterium-mediated transient assays. (**A**) Diagram of the overlapping natural antisense pair *At4g39838* and *At4g39840*, with transcription start sites indicated by arrows and genome coordinates shown. 35S-driven constructs express long (L) and short (S) sequence versions of the *At4g39838*-derived long non-coding RNA. The predicted sequence interacting with the TCA-rich region of the *BIR1* 5’-leader is shown in grey. (**B**) Western blot of HA-tagged BIR1 in leaves co-infiltrated with lncRNA (S or L) and 35S-driven WT BIR1 constructs, with or without the 5’-leader. (**C**) Western blot of HA-tagged BIR1 in leaves co-infiltrated with 35S-driven lncRNAs harboring Arabidopsis miR399, miR158, or miR171 precursors, and WT *BIR1* constructs, with or without the 5’-leader. Controls (-) were infiltrated with empty vector (EV). (**D**) Western blot of HA-tagged BIR1 in leaves co-infiltrated with lncRNA (S or L) and WT or mutated (mA-1) *BIR1* constructs containing the 5’-leader. EV-infiltrated leaves served as controls in (B) and (D). Samples were collected at 2 dpi. Immunoblots were probed with anti-HA antibody; Ponceau staining shows loading. Protein standards (KDa) are indicated. Experiments were independently repeated with consistent results.

### Mutations at or near cleavage site A have no effect on *BIR1* expression

Cleavage site A was consistently identified as a major degradation hotspot in degradome libraries from Arabidopsis leaves and inflorescences (Guzman-Benito *et al*., 2019). This site, along with nearby secondary cleavage sites, maps within a TCA-repeat-rich region of the *BIR1* 5’-leader sequence (Guzman-Benito *et al*., 2019). We further explored the role of the cleavage site A and additional downstream sequences in regulating *BIR1* expression. HA-tagged *BIR1* WT and mA-1 mutant were transiently expressed in *N. benthamiana* under the 35S promoter (**Figure 2A**). Northern and Western blot analyses showed that the mA-1 mutation had little to no effect on *BIR1* mRNA and protein levels (**Figures 2E, F**), consistent with TRV-mediated expression of HA-tagged *BIR1* mA-1 (**Figure 2G**) and with the transient expression of a mCherry-tagged *BIR1* mA-1 mutant under a DEX-inducible promoter (**Figure 3C**). We next generated two additional *BIR1* mutants, an insertion mutant (mA-2) and a deletion mutant (mA-3), targeting sequences downstream of site A, expressed using TRV as a viral vector (**Figure 5F**). Western blot analysis of systemically infected leaves revealed similar BIR1 protein levels in WT and mA-1, with modest reductions in mA-2 and mA-3. As expected, the mBCD control showed higher BIR1 accumulation than WT or any mA variant (**Figure 5G**). These findings suggest that mutations at or near site A have limited effect on *BIR1* expression *in vivo*. Comparable TRV levels across samples confirm that differences in *BIR1* expression were not due to infection variability (**Supplemental Figure 1C**). Interestingly, the repressive effects of the 5’-leader on translation persisted when cleavage site A was mutated (mA-1) (**Figure 5D**). Similarly, repression also occurred when lncRNA was co-expressed with a *BIR1* mA-1 construct, indicating that site A mutations in our study are not sufficient to prevent lncRNA-mediated regulation (**Figure 7D**).

### Identification of potential site A-interacting miRNAs in Arabidopsis

Using the CleaveLand4 pipeline to compare *BIR1* degradome signatures with miRBase, we identified ath-miR5658 as a candidate miRNA potentially mediating cleavage at site A within the 5’-leader (**Supplemental Figure 2C**). The predicted cleavage site corresponds to the tenth nucleotide of ath-miR5658, consistent with miRNA-guided slicing. Partial complementarity to downstream 5’-leader sequences may explain alternative cleavage events observed previously by PARE and 5’-RACE (**Supplemental Figure 2C**) (Guzman-Benito *et al*., 2019). Although ath-miR5658 maps to a genomic region capable of forming a hairpin (**Supplemental Figure 2D**) and was annotated from AGO-bound sRNAs (Axtell *et al*., 2006), we failed to detect its mature or passenger strands by sRNA blotting, stem-loop RT-qPCR, or library sequencing in WT or silencing mutants (data not shown). Additionally, overexpression of the predicted precursor did not yield detectable signals (**Supplemental Figure 2E**), suggesting ath-miR5658 is not a *bona fide* miRNA in Arabidopsis. Interestingly, ath-miR5658 aligns to the natural antisense transcript *At4g39838*, overlapping the stress-responsive *At4g39840* (**Supplemental Figure 3A**). Virus infection increased sense and antisense virus-activated siRNAs (vasiRNAs) at this locus, indicating these sRNAs, including the putative annotated ath-miR5658, derive from dsRNA intermediates rather than canonical DCL1-processed fold-back precursors (**Supplemental Figures 3A**). Further analysis identified another sRNA with near-perfect complementarity to site A and adjacent positions within the TCA-rich region (**Supplemental Figure 2F**), originating from five Arabidopsis genes (**Supplemental Figure 2G**). Among these, *At1g42990* produced abundant vasiRNAs in virus-infected but not mock-treated plants (**Supplemental Figure 3B**). This sRNA locus lacks stem-loop potential, supporting its classification as a vasiRNA. Despite these findings, mutations disrupting sRNA binding at site A had minimal impact on *BIR1* expression, suggesting that cleavage within the 5’-leader is likely regulated by mechanisms independent of sRNA-mediated silencing.

## Discussion

RNA silencing plays a key role in regulating endogenous genes under various stress conditions. For instance, during viral infection, vasiRNAs derived from endogenous genes accumulate and likely guide the silencing of their gene of origin (Cao *et al*., 2014). Degradome analyses across plant species have revealed cleavage patterns consistent with vasiRNA-mediated endonucleolytic activity, although only a small subset of vasiRNAs is actually predicted to mediate cleavage (Pitzalis *et al*., 2020) (unpublished data). In this study, we identified novel candidate RNA silencing targets in Arabidopsis by detecting increased accumulation of cleaved 3’ mRNA products in virus-infected plants compared to mock controls, supporting the idea that viral infection triggers sRNA-guided post-transcriptional regulation of endogenous transcripts. Among them, the receptor-like kinase *BIR1* emerges as a virus-responsive gene undergoing selective degradation.

Previous work and this study showed that *BIR1* transcripts accumulate in RNA silencing mutants during infection, suggesting post-transcriptional silencing helps remove excess *BIR1* transcripts produced during viral activation (Guzman-Benito *et al*., 2019). Consistent with this, *BIR1*-derived siRNAs accumulate widespread in infected but not mock-treated plants (Guzman-Benito *et al*., 2019). Using 5’ RACE, we identified a novel cleavage site (D), complementing two other sites (B and C) previously detected via degradome sequencing (Guzman-Benito *et al*., 2019). These sites represent preferential cleavage hotspots within the *BIR1* coding region. *BIR1* constructs carrying simultaneous mutations at B, C, and D accumulated more mRNA and protein than the WT in both Arabidopsis and *N. benthamiana*, across three expression systems using HA or mCherry C-terminal tags, indicating that these mutations impair transcript cleavage. We found that single-site mutations had, in general, weaker effects, suggesting that all three sites contribute to degradation. WT *BIR1* transcripts normally accumulate at lower levels than the triple mutant (mBCD), but reach similar levels in NbDCL2/4i lines, which are defective in siRNA biogenesis. This supports a model in which *BIR1*-derived vasiRNAs mediate *cis*-silencing through transcript cleavage. However, although *BIR1*-derived vasiRNAs are produced throughout the *BIR1* sequence, our results demonstrate that only a small fraction of the total are capable of guiding the processing of target transcripts and contributing to silencing. Likely, the predominance of specific cleavage sites may reflect sequence or structural preferences of AGO-siRNA complexes (Overhoff *et al*., 2005). Interestingly, post-transcriptional silencing fails to contain *BIR1* expression when the transcriptional regulation is lost and the gene is highly transcribed over time, as in our DEX-inducible system. This suggests that cleavage supports transcriptional regulation functioning as a fine-tuning mechanism to maintain *BIR1* transcript levels within a functional threshold. Notably, cleavage sites B and C are located 13 and 40 nucleotides upstream of the stop codon, respectively, a region often associated with ribosome pausing and co-translational decay (Hou *et al*., 2016; Yu *et al*., 2016). The 28-nt spacing between B and C matches the footprint of a stalled ribosome (Pelechano *et al*., 2015), raising the possibility that ribosome pausing near the stop codon may contribute to the observed cleavage patterns. However, this model does not fully explain why synonymous mutations at these sites increase *BIR1* transcript and protein levels, even when using HA- or mCherry-tagged versions. Alternative explanations, such as sequence-specific cleavage by other endonucleases in infected tissue, cannot be ruled out.

Along with RNA silencing, sRNA-independent mechanisms may also contribute to processing. Matching *BIR1* degradome data with miRBase revealed that the 5’ degradome signature at site A aligns with the predicted cleavage site of ath-miR5658 (Guzman-Benito *et al*., 2019). Additional nearby signatures detected by PARE and 5’ RACE in this former study also showed partial complementarity to ath-miR5658, indicating potential suboptimal cleavage events (Guzman-Benito *et al*., 2019). However, although ath-miR5658 is listed in miRBase, it may not meet established plant miRNA criteria. Structural prediction places ath-miR5658 in a fold-back region (ΔG = -36.90 kcal/mol), but it is rarely detected in deep sequencing, and no miRNA* strand has been confirmed. In addition to ath-miR5658, another vasiRNA was found to exhibit near-perfect complementarity to the 5’ leader of *BIR1* at cleavage site A and adjacent positions. This siRNA aligns with five Arabidopsis genes, including *BASIC REGION/LEUCINE ZIPPER MOTIF 60* (*bZIP60*), a source of abundant vasiRNAs during viral infection. These findings suggest that *BIR1* silencing may be mediated by a network of siRNAs, rather than a single trigger. Nevertheless, unlike mutations at sites B, C, and D, nucleotide changes at or near cleavage site A do not affect *BIR1* transcript or protein levels, suggesting that degradation at site A may not rely on single sequence-specific recognition or DCL2/4-dependent siRNA/vasiRNA pathways. Conversely, our results suggest that cleavage at site A is not a regulatory event *per se*, but rather a downstream consequence of inhibited BIR1 protein synthesis. Additional studies are needed to clarify this hypothesis.

A key finding of our study is the role of the *BIR1* 5’ leader in regulating translation. We found that the 5’ leader acts as a repressive element, in which both linear and structural features may be important translational checkpoints (Hinnebusch *et al*., 2016; Leppek *et al*., 2018). We identified six type 1 uORFs located entirely within the 5’ leader, of which uORFs 4, 5 and 6 are translated. These *cis*-regulatory elements can interfere with ribosome scanning and suppress downstream mORF translation via ribosome stalling, leaky scanning, or limited reinitiation (Kurihara *et al*., 2018; von Arnim *et al*., 2014; Wu *et al*., 2022). In plants, uORF-mediated translational repression has been extensively described for many immune and stress-associated mRNAs both under normal conditions and upon pathogen attack or stress (Nazarian-Firouzabadi *et al*., 2025). Here, analysis of Ribo-seq data revealed active translation of the *BIR1* uORFs under both mock and ETI conditions in transgenic Arabidopsis plants expressing *DEX::AvrRpt2* (Zhou *et al*., 2023), indicating their potential regulatory role. One possibility is that these uORFs recruit ribosomes to initiate upstream translation, after which uORF-associated ribosomes reinitiate at the downstream main ORF. Such a mechanism would lead to proportional increases in ribosome occupancy on both uORFs and the mORF upon ETI induction. Given that these uORFs are efficiently recognized and translated by ribosomes, we also cannot exclude an alternative mechanism in which they function through their encoded peptides. In this scenario, uORF-derived peptides could act *in trans* to influence *BIR1* translation or signaling, consistent with their demonstrated translational competence (Mou *et al*., 2025). lncRNAs are known to regulate translation by interacting with RNA elements within the 5’ UTR (Leppek *et al*., 2018). We identified a *At4g39838*-derived lncRNA that modulates *BIR1* translation efficiency. This effect was dependent on the 5’ leader region, and was mediated by sequence complementarity, as other known lncRNAs (carrying miRNA precursors) had no impact on *BIR1* expression. Accordingly, we propose that the TCA-rich region within the 5’ leader of *BIR1* serves as a complementary binding site for this lncRNA. The precise mechanism of action remains unclear, but the lncRNA may interfere with other regulatory motifs within this region or alter mRNA stability.

In summary, our results contribute to advancing the understanding of the mechanisms regulating *BIR1* expression, which may be shared by other members of the gene family. These findings confirm the role of RNA silencing in the regulation of *BIR1*, delineating the sequences susceptible to sRNA-mediated cleavage. Although further research is needed, our data highlight the importance of the 5’ leader in regulating BIR1 protein levels, likely through regulatory elements such as uORFs and lncRNA.

## Acknowledgments

The funders had no role in study design, data collection and analysis, decision to publish, or preparation of the manuscript.

## Author contributions

C.L. and C.M. conceived and designed the study; I.G.B., C.R., L.D., L.F.C., J.M.F.-Z., and C.L. outlined experiments, performed research and analysed data; J.M.F.-Z., L.D, C.M., R.N. and C.L. contributed with bioinformatic analysis; R.N. and G.X. provided Riboseq data; C.L. wrote the paper.

## Supplementary data

**Table S1:** List of primers

**Table S2:** Potential RNA silencing target genes detected in mock-inoculated and TRV-infected inflorescences

**Supplementary Figure 1.** Mutations at predicted cleavage sites within the *BIR1* coding sequence enhance transcript accumulation.

**Supplementary Figure 2**. Characterization of regulatory elements within the 5’-leader of *BIR1*.

**Supplementary Figure 3.** Identification of *BIR1*-interacting virus-associated small RNAs in Arabidopsis.

**Supplementary Figure 1.**
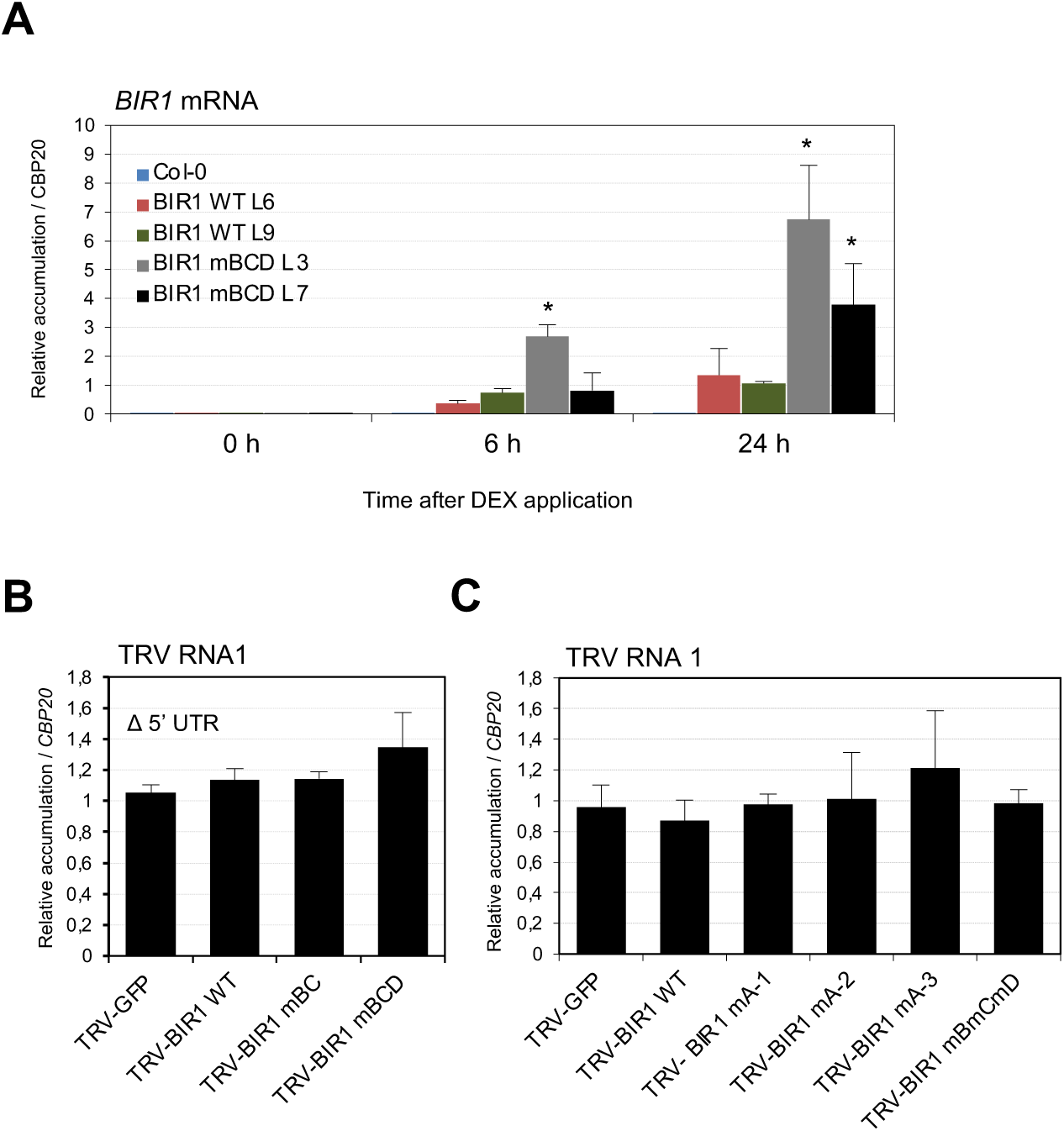
Mutations at predicted cleavage sites within the *BIR1* coding sequence enhance transcript accumulation. (**A**) RT-qPCR analysis of *BIR1-mCherry* transcripts in DEX-inducible transgenic Arabidopsis lines expressing wild-type (WT: L6, L9) or mutated (mBCD: L3, L7) *BIR1* after 6 and 24 h of DEX treatment (30 µM). (**B**) RT-qPCR of TRV RNA 1 in upper leaves at 5 dpi, systemically infected with TRV derivatives expressing GFP or BIR1 variants (WT, mBC, mBCD) lacking the 5’-leader (Δ5’ UTR). (**C**) RT-qPCR of TRV RNA 1 in upper leaves at 5 dpi, infected with TRV derivatives expressing GFP or BIR1 variants (WT, mA-1, mA-2, mA-3, mBCD). All RT-qPCR data were normalized to *CBP20*.

**Supplementary Figure 2.**
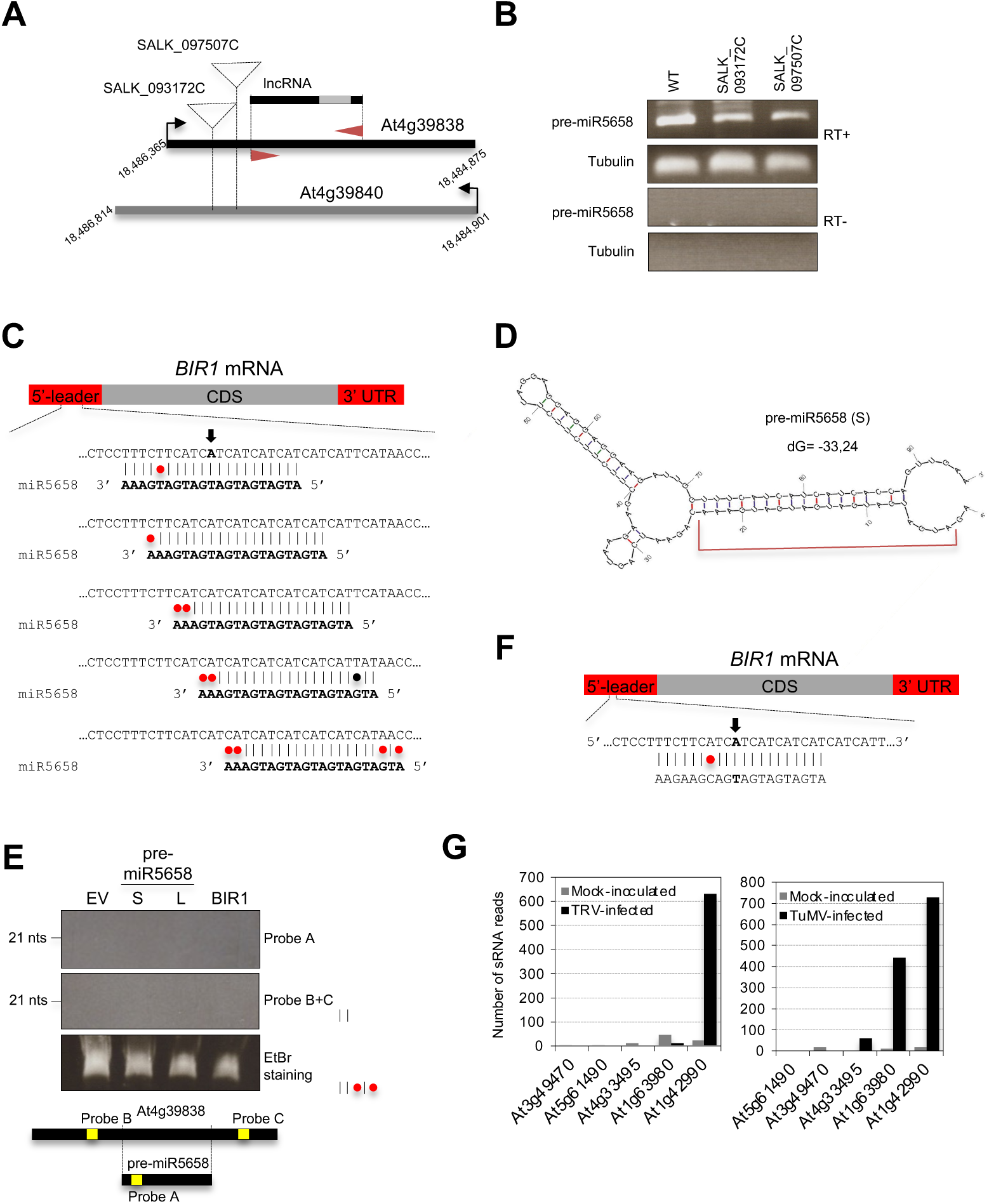
Characterization of regulatory elements within the 5’-leader of *BIR1*. (**A**) Schematic of the *At4g39838* locus and T-DNA insertion sites (SALK_097507C, SALK_093172C) shown as dotted triangles; transcription start sites as black arrows. (**B**) RT-PCR of lncRNA transcripts derived from the *At4g39838* locus in WT and T-DNA lines, with (+) or without (–) reverse transcriptase. β-Tubulin was used as a control. (**C**) Predicted ath-miR5658 interaction sites within the TCA-rich 5’ untranslated leader region of *BIR1*. G-U base pairs (black dots) and mismatches (red dots) are indicated. The major cleavage site A at position 156 (from degradome sequencing) is marked with a thick arrow. (**D**) Predicted secondary structure of the pre-miR5658 RNA. The mature ath-miR5658 sequence is highlighted within the stem-loop. **(E)** RNA blot of small RNAs from leaves infiltrated with 35S-driven pre-miR5658 constructs at 2 days post-infiltration (dpi). Blots were probed with A (ath-miR5658-complementary) and B/C (outside the pre-miR5658 region) as shown. Empty vector (EV) was used as control. A 21-nt size marker is indicated. **(F)** Predicted interaction between a BIR1-targeting small RNA and the TCA-rich 5’ leader. (**G**) Arabidopsis loci matching the candidate BIR1-interacting sRNA and virus-associated siRNA (vasiRNA) production in mock- and virus-infected plants (tobacco rattle virus (TRV), left; turnip mosaic virus (TuMV), right).

**Supplementary Figure 3.**
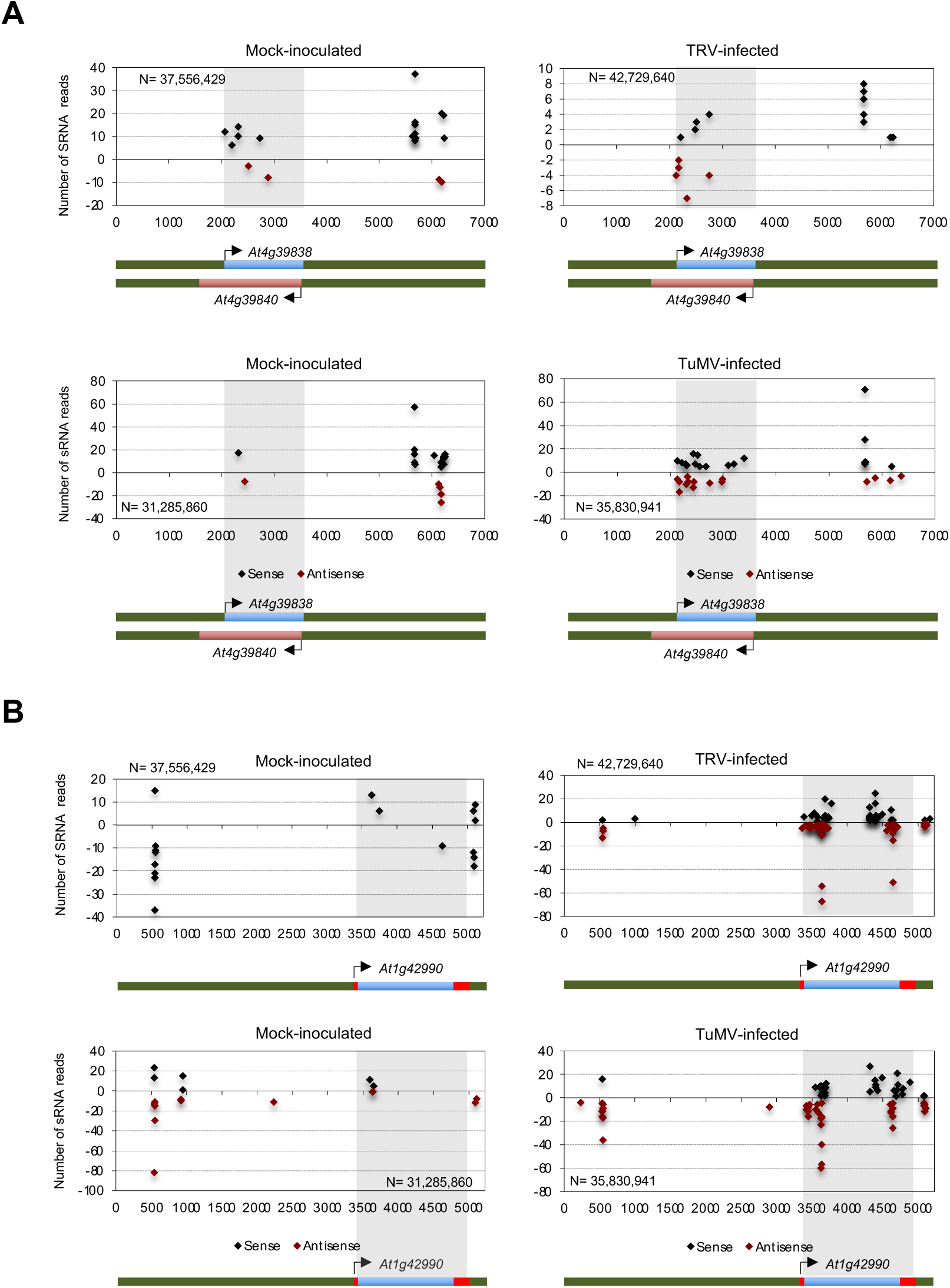
Identification of *BIR1*-interacting virus-associated small RNAs in Arabidopsis. (**A**) Distribution of small RNAs (sRNAs) derived from *At4g39838* in mock- and turnip mosaic virus (TuMV)- or tobacco rattle virus (TRV)-infected Arabidopsis plants. The shaded region marks the natural antisense overlap between *At4g39838* and *At4g39840*. (**B**) Distribution of sRNAs derived from *At1g42990* in mock- and TuMV- or TRV-infected plants. The shaded region represents the *At1g42990* coding sequence. Sense (black dots) and antisense (red dots) sRNAs are plotted as positive and negative values, respectively. N indicates the total number of filtered small RNA reads.

